# The activation chain of the broad-spectrum antiviral bemnifosbuvir at atomic resolution

**DOI:** 10.1101/2024.02.16.580631

**Authors:** Aurélie Chazot, Claire Zimberger, Mikael Feracci, Adel Moussa, Steven Good, Jean-Pierre Sommadossi, Karine Alvarez, François Ferron, Bruno Canard

**Affiliations:** AFMB, CNRS, Aix-Marseille University, UMR 7257, Case 925, 163 Avenue de Luminy, 13288 Marseille Cedex 09, France; ATEA Pharmaceuticals, Inc.; 225 Franklin St., Suite 2100, Boston, MA 02110 USA

## Abstract

Bemnifosbuvir (AT-527) and AT-752 are guanosine analogues currently in clinical trials against several RNA viruses. Here we show that these drugs require a minimal set of 5 cellular enzymes for activation to their common 5’-triphosphate AT-9010, with an obligate order of reactions. AT-9010 selectively inhibits essential viral enzymes, accounting for broad spectrum antiviral potency. Functional and structural data at atomic resolution decipher *N^6^*-purine deamination compatible with metabolic activation by human ADALP1. Crystal structures of human HINT1, ADALP1, GUK1, and NDPK at 2.09, 2.44, 1.76, and 1.9 Å resolution, respectively, with cognate precursors of AT-9010 illuminate the activation pathway from the orally available bemnifosbuvir to AT-9010, pointing to key drug-protein contacts along the activation pathway. Our work provides a framework to integrate the design of antiviral nucleotide analogues, confronting requirements and constraints associated with activation enzymes along the 5’-triphosphate assembly line.

## Introduction

The recent COVID19 crisis due to severe acute respiratory syndrome coronavirus 2 (SARS-CoV-2) has evidenced the need for safe and potent antivirals, in conjunction with accurate and rapid diagnostics. First, at the public level, prophylaxis around index cases may significantly curb emergence by cutting viral load and transmissibility. Second, at the individual level, early therapy may limit virus spread to vital organs, avoiding complications.

Nucleoside/nucleotide analogues (NAs) represent the first family of compounds which have been used as direct-acting antivirals several decades ago^1,2,3^ against DNA viruses such as herpesvirus, hepatitis B virus (HBV), and human immunodeficiency virus type 1 (HIV-1), and RNA viruses such as hepatitis C virus (HCV) and SARS-CoV-2. Upon reaching their target cells, antiviral drugs often act after intracellular activation. Early biologically active nucleoside analogues (NAs) such as ara-C or cytarabine penetrate the target cell efficiently and are transformed into the nucleoside analogue 5’-triphosphate (NA-TP) by a series of kinase reactions (reviewed in^2–4^).

ProTide prodrug technology has been implemented to deliver NA monophosphates^5^. Aryloxy phosphoramidate prodrugs of *eg.*, tenofovir disoproxil fumarate (TDF) for HIV-1 and HBV, sofosbuvir (SOF) for HCV, and remdesivir (RDV) for SARS-CoV-2 have met clinical success^5–7^. They achieve cell penetration through shielding the phosphate charges, and preferential hydrolysis at the P-N bond by-passes the first -often limiting-nucleoside kinase reaction, allowing significant building up of NA-TP pools. NA-TPs poison viral RNA synthesis specifically, provoking premature viral RNA chain termination or chemical/genetic corruption of the viral nucleic acid^8^.

Two main parameters determine NA potency: the concentration ratio of NA-TP over its natural NTP competitor, and the selectivity of the viral RdRp for use of the NA-TP^9^. NA-TP pools must be formed efficiently upon intracellular activation, and the differences of the NA-TP scaffold relative to its natural NTP counterpart must remain ‘below the radar’ of the viral RdRp.

General knowledge, structural and functional data about cellular enzymes along a given NA activation pathway are fragmented. Numerous NAs have been designed showing appropriate poisoning of viral RNA synthesis through their 5’-TP *in vitro*, but clinical development has failed for lack of transmembrane permeability, intracellular metabolic activation, and/or cellular toxicity. Clearly, nucleotide analogue drug-design needs an integrated view (*ie.*, structural, functional, and mechanistic) from the delivered NA up to the ultimate inhibited viral enzyme reaction accounting for antiviral effect, as well as the molecular basis for (non)interaction with cellular enzymes.

Such a global picture is emerging for two FDA-approved NA prodrugs directed against RNA viruses, SOF and RDV (Fig. S1). The activation pathway of RDV has been elucidated and profiled in a variety of tissues^10^. RDV is an aryloxy phosphoramidate prodrug (GS-5734, Veklury) converted in two steps to the 5’-monophosphate NA by esterases belonging to two families (CatA and CES1), followed by the cleavage of the P-N bond by a histidine triad nucleotide (HINT) phosphoramidase^11^. The resulting monophosphate NA is then converted to RDV-TP by subsequent action of two cellular phosphotransferases. However, although detailed studies at the atomic level exist to understand accommodation of the 1’-CN group at the SARS-CoV-2 RdRp active site^12,13^, structural insight of the relevant activation intermediates is lacking for all enzymes in the activation pathway.

SOF (PSI-7851, Sovaldi) undergoes the same CatA/CES1 deprotection pathway as RDV, but is activated to the 5’-triphosphate by the UMP-CMP kinase and the nucleoside diphosphate kinase (NDPK) sequentially^14^. Here again, although the mechanism of chain-termination at the HCV NS5b RdRp has been elucidated at atomic resolution^15^, the structural basis of activation remains uncharacterized.

Bemnifosbuvir (AT-527 (hemisulfate salt), AT-511 (free base)) is a guanosine analogue currently in clinical trials against SARS-CoV-2 and HCV. It showed in Phase II clinical trials a 71% risk reduction in out-patients with moderate COVID-19 (MORNINGSKY; NCT04396106)^16,17^, and is currently under investigation in a global phase III clinical trial in outpatients at high risk for disease progression (SUNRISE-3; NCT05629962). It is also being evaluated as an anti-HCV drug^18^ in combination with the NS5A inhibitor ruzasvir (NCT05904470). Its epimer AT-752 is currently in clinical phase II against Dengue virus (NCT05466240)^19,20^.

Both bemnifosbuvir and AT-752, once processed by the CatA/CES1 pathway, converge to the same precursor AT-551 (Fig. 1A)^18^. These two analogues are amongst the few antiviral purine nucleotide analogues devoid of significant cellular toxicity. Once the P-N bond of AT-551 is hydrolyzed (presumably) by HINT1^21^, giving rise to the monophosphate AT-8003, the diamino purine base is (presumably) converted to a natural guanosine base through specific *N^6^*-deamination carried out by the ADALP1 enzyme^22,23^. This reaction is believed to skirt a cellular step responsible for the toxicity of unprotected, ‘natural’ guanosine analogues^18,24^. The resulting 2’-F-2’-C-methyl guanosine 5’-monophosphate (AT-8001) is (presumably) consecutively phosphorylated twice to yield AT-9010 by means of guanylate kinase 1 (GUK1)^25^ and nucleotide diphosphate kinase (NDPK)^26,27^. AT-9010 accumulates in various cell types^18^. The HCV RNA synthesis is likely halted through RNA chain termination^18^. SARS-CoV-2 RNA synthesis is halted through dual targeting of the replicase complex^28^, and the Dengue virus RNA synthesis is halted through dual targeting of the viral protein NS5^19,20^.

**Figure 1.**
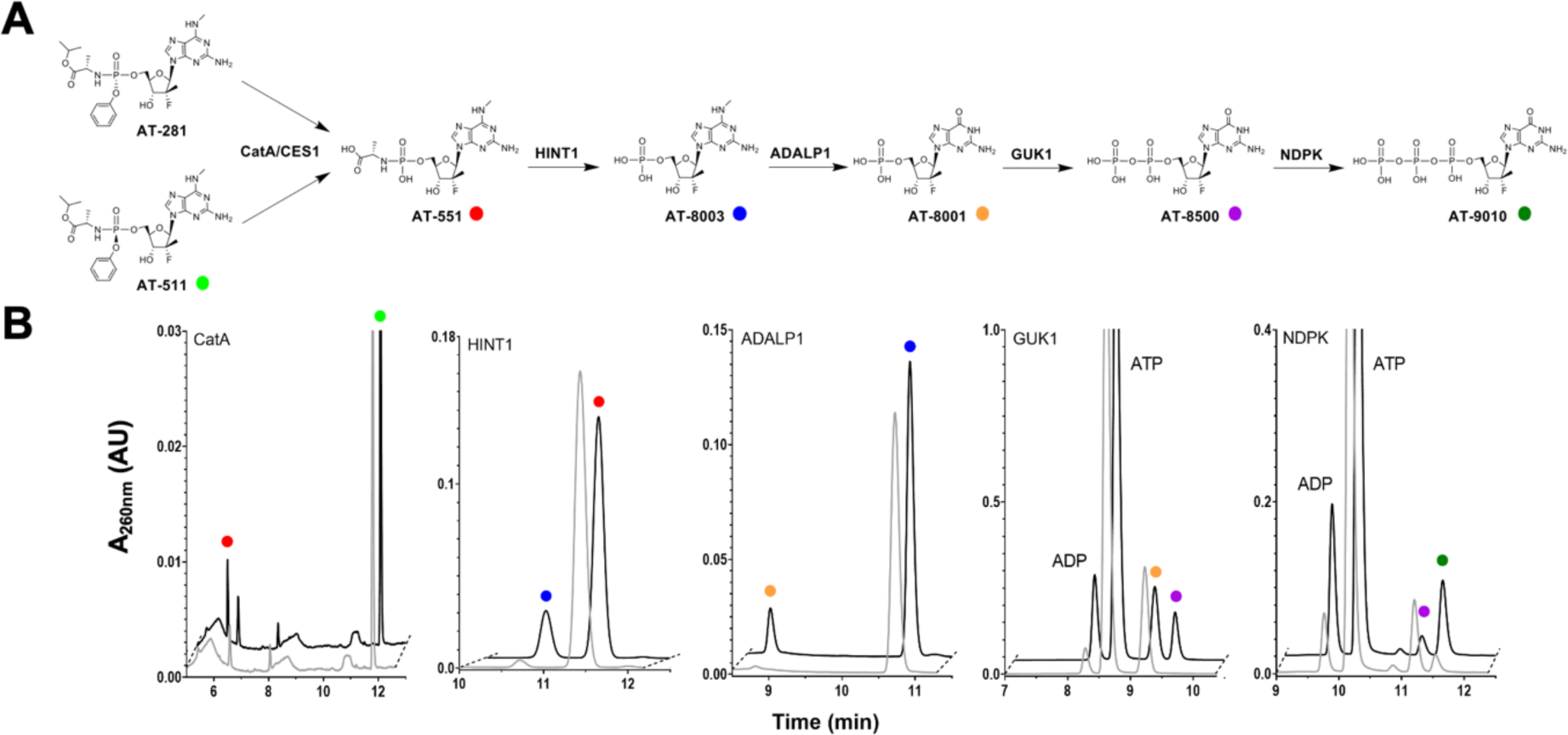
Validation of the activation pathway of bemnifosbuvir. (A) The activation pathway of AT-511, the free base form of bemnifosbuvir (AT-527), into the active form AT-9010. AT-281, the diastereoisomer of AT-511 and free base form of AT-752 follows the same pathway. (B) Typical RP-HPLC chromatograms of the enzymatic assays after 2 min (grey) and the last kinetic point (black). For each enzyme CatA, CES1, HINT1, ADALP1, GUK1 and NDPK, the successful validation of activity results in conversion of the substrate into the corresponding product. All compounds were identified by co-injection with authentic samples.

Most information relative to activation pathways has been obtained through measurements of intracellular concentrations of NA intermediates^10,11,14,24^. Studies described herein aim to ascertain which individual enzymes are involved in the activation pathway, as well as clarify their structural and functional mode of interaction with activation intermediates. Interaction maps of NAs with both activating partners and viral enzymes at atomic resolution would guide improved design of novel NAs.

In this work, we have identified a minimal set of 5 human enzymes involved in the activation of bemnifosbuvir and AT-752 to their active 5’-TP form AT-9010. We demonstrate distinct stereo-preference of CatA and CES1 for AT-527 and its epimer AT-752, respectively. We elucidate an ordered activation pathway with potential limiting steps, including substrate specificity requirements at the *N^6^*-purine position for ADALP1. Structural analysis with key human enzymes HINT1, ADALP1, GUK1, and NDPK in complex with the bemnifosbuvir/AT-752 intermediate NA at 2.09, 2.44, 1.76, and 1.9 Å resolution informs on both the *N^6^*-purine and 2’-ribose chemo-steric requirements leading to active NAs targeting viral RdRps.

## Results

The putative metabolic pathway of AT-527 (bemnifosbuvir) and AT-752 is shown in Fig. 1A.

Cathepsin A (CatA) and/or carboxylesterase 1 (CES1) presumably precede a spontaneous decomposition yielding the L-alanine phosphoramidate prodrug AT-551, common to both AT-752 and AT-527 pathways. AT-551 is then subjected to four enzyme-mediated reactions leading to the active 5’-triphosphate form, AT-9010. Apart from the *N^6^*-deamination reaction, this pathway has been inferred from that of the anti-HCV drug SOF, which carries a uracil base instead of the *N^6^*-methyl diamino purine base (a guanine precursor) of either AT-752 or AT-527. The SOF pathway, however, has been determined indirectly, by measuring intermediates in cells treated with SOF^11,14^.

### Five enzymes are needed to convert AT-527 (bemnifosbuvir) or AT-752 to their active form AT-9010

We expressed, purified and crystallized four of the human enzymes widely distributed in human tissues (https://proteinatlas.org) and supposedly involved in the prodrug activation pathway (Fig. 1A, Fig. S2). Together with human enzymes CatA and CES1, we challenged HINT1, ADALP1, GUK1, and NDPKb with their anticipated substrates depicted in Fig. 1A and Fig. S3. We made use of an HPLC-UV method to follow the conversion of the substrate, together with a kinetic monitoring relative to known standards (Fig. 1B).

Both AT-527 and AT-752 can be enzymatically converted to AT-9010 by an ordered series of reaction involving non-specific esterases CatA/CES1, HINT1, ADALP1, GUK1, and NDPK (Table 1). For AT-511, non-specific esterases CatA and CES1 exhibit a ∼300-fold difference in activity whereas hydrolysis efficiency is quite similar for AT-281 (< 3-fold difference). Both for control and AT compounds, the early steps of the pathway are the slowest, with di- and triphosphate formation showing the fastest turnovers.

**Table 1:** Activities of enzymes involved in AT-527/AT-511 and AT-752/AT-281 activation.

| Enzyme | Substrate | Specific activity<br>(nmol.min<sup>-1</sup>.nmol<sup>-1</sup>)<sup>a</sup> |
| --- | --- | --- |
| CatA | **AT-511** | 9.8 ± 0.3 |
|  | **AT-281** | 0.7 ± 0.1 |
| CES1 | **AT-511** | 0.033 ± 0.004 |
|  | **AT-281** | 0.27 ± 0.02 |
| HINT1 | AMP-NH2 | 101 ± 8 |
|  | **AT-551** | 0.7 ± 0.2 |
| ADALP1 | N<sup>6</sup>-Me-AMP | 42 ± 2 |
|  | **AT-8003** | 9.4 ± 0.4 |
|  | AT-8010 | 5.21 ± 0.07 |
|  | AT-8004 | 11.9 ± 0.1 |
|  | AT-8002 | 0.94 ± 0.03 |
|  | AT-551 | No activity |
|  | AT-229 | No activity |
|  | AT-259 | No activity |
| GUK1 | GMP | 6800 ± 300 |
|  | **AT-8001** | 34 ± 5 |
|  | AT-8003 | No activity |
| NDPKB | GDP | 5400 ± 800 |
|  | **AT-8500** | 1300 ± 100 |

<sup>a</sup> Results were obtained from at least 3 independent experiments and reported as means ± standard deviations

### The phosphate stereo-selectivity of CatA/CES1 usage

The phosphorus atom of the aryloxy phosphoramidate moiety is chiral, resulting in two possible epimers. AT-511 (*S*_P_ isomer) is the free base form of AT-527, and AT-281 (*R*_P_ isomer) is the free base form of AT-752 (Fig. S3). As shown for SOF, the difference of antiviral activity between the two stereoisomers could be due to the activation of the prodrug and more specifically to the stereo-selectivity of the enzymes involved in the first hydrolytic step.

CatA or CES1 were incubated with either AT-511 or AT-281 and comparative velocities of AT-551 formation were measured (Fig. S4). CatA always shows higher activity than CES1. CatA prefers by 14-fold the *S*_P_ isomer AT-511 over the *R*_P_ isomer AT-281 (9.8 ± 0.3 *vs*. 0.7 ± 0.1 nmol/min/nmol protein). Interestingly, however, a reversed stereo-selectivity is observed for CES1 relative to CatA: CES1 hydrolyses AT-281 ∼10-fold faster than AT-511 (0.27 ± 0.02 vs. 0.033 ± 0.004 nmol/min/nmol protein, respectively). These results align with those obtained with a variety of clinically relevant prodrugs; for SOF, TAF, and RDV, CES preferentially hydrolyzes the *R*_P_ isomer whereas CatA prefers the *S*_P_ isomer of the aryloxy phosphoramidate moiety^10,11,14^. We conclude that non-specific esterases of the CatA/CES1 type are able to convert *bona fide* McGuigan ProTides into HINT1 substrates from both AT-752 and AT-527. The respective abundance of each enzyme in cells or tissues might be useful as a predictor of ester hydrolysis efficiency in these given cells or tissues.

### Specificity and order of the reactions

AT-551 could not act as a substrate for ADALP1 (Fig. S5). This result indicates that the L-alanine phosphoramidate precludes efficient either binding or catalytic conversion of AT-551 with ADALP1 (see structural data below). AT-8003 could neither be a substrate for GUK1, indicating that the *N^6^* position must be either unsubstituted or only an O^6^ to allow binding and phosphorylation of the NMP (Fig. S5). Unless other enzymes exist that could use these substrates and by-pass the proposed activation sequence, the absence of reactivity reported here is consistent with the order of the reaction pathway reported in Fig. 1A.

### Structural and functional analysis of HINT1 substrate specificity

HINT1 belongs to the HIT (histidine triad) protein superfamily, and has been shown to hydrolyse the P-N bonds in SOF and RDV^10,11,14^. HINT1 is able to hydrolyze AMP-NH_2_ (control) into AMP, and AT-551 into AT-8003 with specific activities of 101 ± 8 and 0.7 ± 0.2 nmol/min/nmol protein, respectively (Fig. 1B and Table 1). HINT1 is thus ∼140-fold faster with the control compound than with AT-551.

We crystallized HINT1 in complex with AT-551 or AT-8003 (Fig. 2).

**Figure 2.**
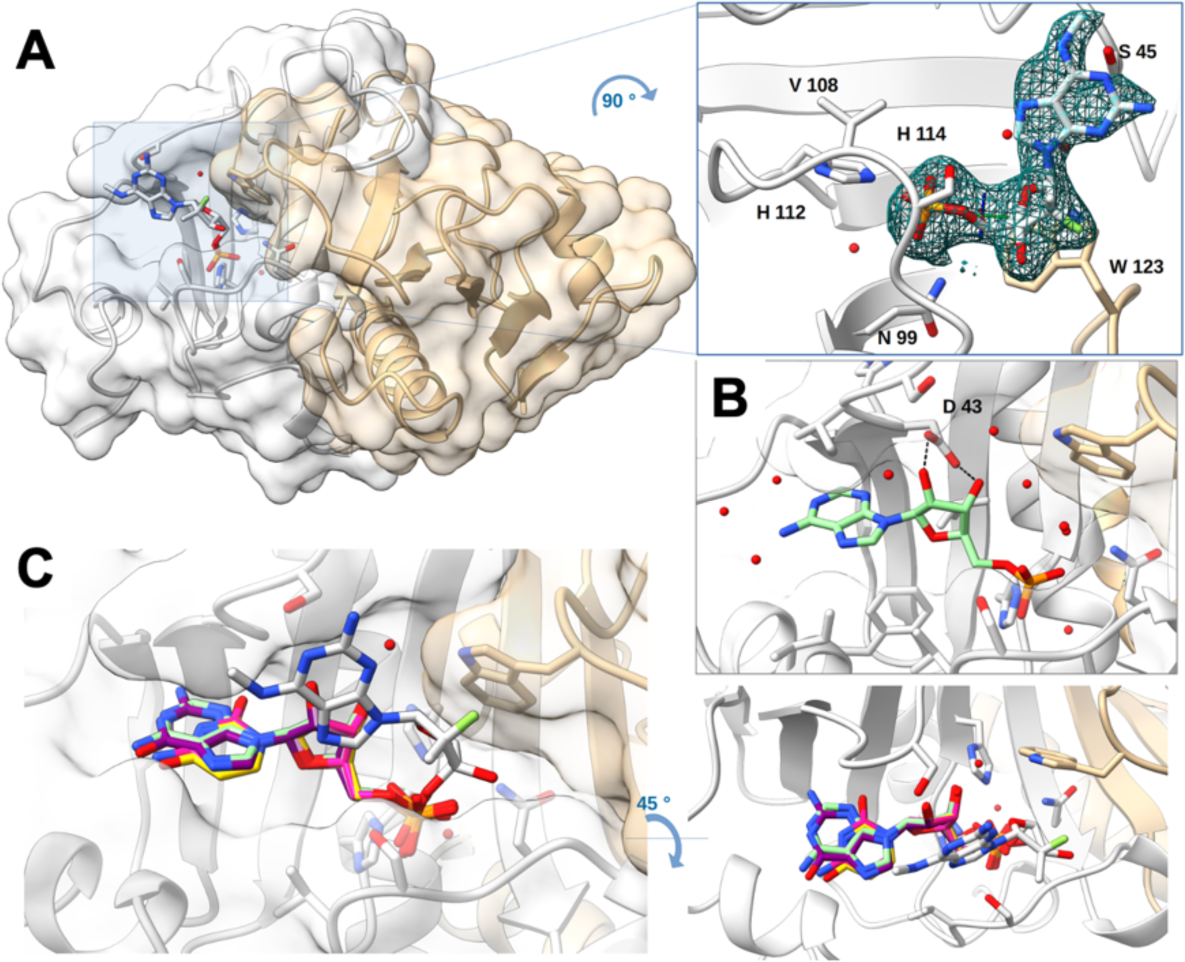
Structure of human HINT1 in complex with AT-8003. **A)** Homodimer of human HINT1 structure complexed with one AT-8003 compound. The protein structure is represented in ribbon and transparent surface, while compound is represented in sticks with heteroatom colors. On the right side an enlargement of the molecule in the active site is presented at 90”. Side chains of interacting residues with the molecule are shown in sticks. The *F_o_ - F_c_* omit map of the molecule is presented at 1 σ. **B)** Binding mode involving direct coordination of the 2’ and 3’ O ribose of AMP (light green - PDB: 5ED6) by Asp 43. **C)** Comparison of the relative positions of AMP (light green - PDB: 5ED6), CMP (orange - PDB: 5KM2), UMP (yellow - PDB: 5KM3), GMP (purple - PDB: 5KM1) in respect to AT-8003 (white) in the HINT1 active site. AT-8003 is positioned almost orthogonally to other nucleotides bound in other structural models, and only the phosphate positions are undistinguishable. The right panel is the same figure turned 45°.

For AT-551, reaction occurred in the crystal resulting in AT-8003 bound to the active site on one monomer (Fig. 2A). The crystal structure (2.09 Å resolution) shows a homodimer, each monomer being almost perfectly superimposable (RMSD ∼0.18 Å on the visible 126 aa).

The bound product pose is significantly different from all existing HINT1 structures^21^ in complex with AMP, GMP, or NMP-phosphoramidate ligands (Fig. 2B). In all available structures so far, the nucleotide analogue is buried in a pocket, contacting conserved amino acids Ile 44 and Asp 43 involved in purine base stacking and ribose 2’,3’-OH ribose bi-dentate binding, respectively. Here, the methylated diamino purine base is bound outside the nucleobase binding pocket, shifted by 90° and making a single hydrogen bond with Ser 45 that could occur with any purine or pyrimidine base. Remarkably, the two parts of the AT-8003 which differentiate it from guanosine are not engaged in any amino acid contact. Both the *N^6^*- amino purine and the 2’-C-methyl, 2’-F are projecting into the solvent.

The phosphate is closely superimposable to that observed in other published structures^21^. The catalytic His 112 nucleophilic nitrogen is 3.1 Å away from the phosphate, and the hydrolytic water molecule is positioned opposite to the putative leaving amine and general acid His 114 and/or His 51. In a similar complex (PDB: 5IPE), the O^6^-guanine of GMP is located at 3.3 Å from the Ile 18 side chain, and there is space to accommodate an *N^6^*-methyl amino group at the solvent-protein interface. The AT-8003 ribose pucker is of *Northern* (*N*) configuration: C3’ up and C2’ down (the *Southern* (S) pucker has the C3’ down and C2’ up), as this conformation must be imposed by the presence of its 2’-C-methyl. The ribose pucker of GMP and AMP analogues in all other structures being *S*, we surmise the 2’-C-methyl is driving this alternate preferential NA binding mode seen in all structures reported here.

### Structural and functional analysis of ADALP1 substrate specificity

ADALP1 is a deaminase acting on *N^6^*-substituted purine nucleoside monophosphates^23^. The activity of purified ADALP1 was tested on *N^6^*-Me-AMP as a control compound before testing on the anticipated substrate AT-8003. Because the anti-HIV drug abacavir bears an *N^6^*- cyclopropyl modification^22^, we tested several *N^6^*-modified analogues: -NH(nPr) (AT-8010), - N(CH_3_)_2_ (AT-8004) and –NH_2_ (AT-8002). The nucleoside versions of AT-8003 and AT-8010 were also tested, *ie.,* AT-229 and AT-259, respectively.

ADALP1 is able to convert *N^6^*-Me-AMP into IMP (42 ± 2 nmol/min/nmol protein), as well as AT-8003, AT-8004 and AT-8010 into AT-8001. AT-8003, AT-8004 and AT-8010 are converted 3 to 8-fold slower than *N^6^*-Me-AMP (9.4 ± 0.4 nmol/min/nmol protein, 11.9 ± 0.1 nmol/min/nmol protein and 5.21 ± 0.07 nmol/min/nmol protein, respectively, Fig. 1B and Table 1). ADALP1 is thus able to accept the 2’-C-methyl, 2’-F modifications on the ribose part, as well as several substituents on the *N^6^*-position. Extension of the *N^6^*-substituent, not its bulk immediately close to the nitrogen atom, seems to negatively influence activity (Fig. 3A).

**Figure 3.**
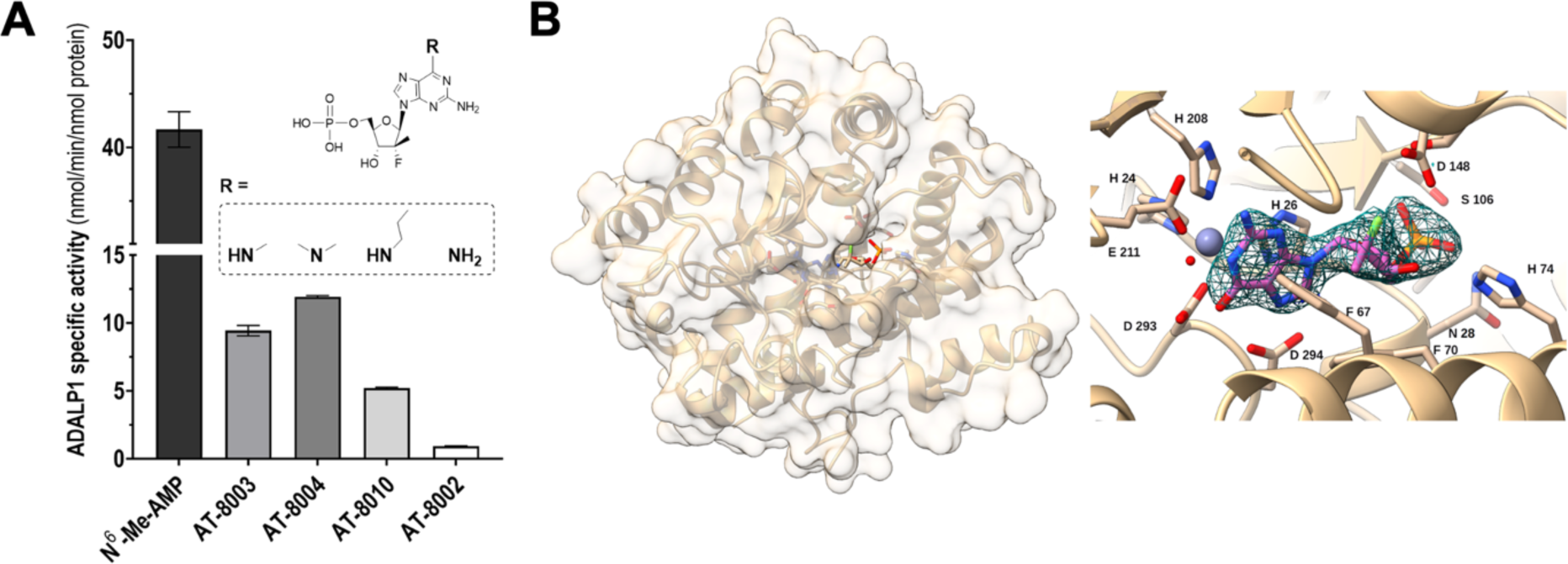
Human ADALP1 analysis. **A)** ADALP1 specificity. Activity of ADALP1 with *N^6^*-Me-AMP as control substrate, AT-8003 as anticipated substrate and *N^6^*-modifíed analogues of AT-8003 : AT-8004 with a *N^6^*-dimethyl group, AT-8010 with a *N^6^*-n-propyl group and AT-8002 with a *N^6^*-amino group. 100nm ADALP1 (or lµM for AT-8002) was incubated 2h at 37°C with 200µM substrate. Bars show mean values (± SD) of three independent experiments. **B)** ADALP1 in complex with AT-8001. Human ADALP1 structure is represented in ribbon and transparent surface, while compound is represented in sticks with heteroatom colors. The molecule is deeply buried in a closed conformation of ADALP1. On the right side a zoomed part of the molecule as well as the zinc ion (grey ball) is presented in the active site. Side chains of interacting residues with the molecule are shown in sticks. The *F_o_ - F_c_* omit map of the molecule is presented at 1 σ.

Under our experimental conditions, ADALP1 converts only nucleoside monophosphate substrates: no activity was detected with the nucleosides AT-229 and AT-259.

Interestingly, the reaction is ∼45-fold slower with AT-8002 as substrate compared to the natural substrate *N^6^*-Me-AMP, indicating that at least one substituent on the nitrogen atom may enhance the leaving group ability.

We co-crystallized ADALP1 and AT-8003. The solved ADALP1 structure adopts an α/β- barrel architecture similar to that of ADAs^29^, with the essential Zn^2+^ ion tightly bound in the vicinity of the catalytic center (Fig. 3B and Fig. S6A). Fourteen α-helices surrounding eight parallel β-strands constitute a TIM-barrel fold^30^. The asymmetric unit contains eight copies of the protein. Since ADALP1 was purified as a homogeneous monomer (Fig. S6B), the octamer likely appeared during crystallogenesis. Although ADALP1 was co-crystallized with the substrate AT-8003, the product AT-8001 was found bound at the active site. The compound is found in all eight chains and all the chains found in the asymmetric unit closely superimpose (r.m.s.d. of ∼0.26 Å) except for the minor loop and N- and C-terminal ends (Fig. S6C). The structural zinc is coordinated by side chains of four residues (His 24, His 26, His 208 and Asp 293) and a water molecule, in a trigonal bipyramidal geometry^31^. Water bridges His 232 and the resulting O^6^ of the purine base suggesting that both His 293 and the water molecule did play a role in the hydrolysis of the amine. AT-8001 is buried in a closed hydrophobic pocket that can only accommodate a monophosphate compound checked and stabilized by Asn 28, His 74, Ser 106, and Thr 107.

Unlike in the HINT1:AT-8003 complex, there is no ambiguity in substrate nor product binding pose^30^. Once the substrate is bound, the catalytic water attacks the purine C^6^ which likely stays at this location whilst the methylamine is leaving in the opposite exit channel.

### Structural and functional analysis of GUK1 specificity

GUK1 is a nucleoside monophosphate kinase able to transfer a phosphate group from ATP to a guanosine 5’-monophosphate, yielding GDP and ADP^25^. Simultaneous binding of monophosphate substrate and triphosphate donor precedes conformational closure and phosphate transfer. This enzyme metabolizes several purine analogues such as abacavir and ganciclovir^32,33^. Expectedly, GUK1 is able to phosphorylate GMP into GDP, but also AT-8001 into AT-8500 (Fig. 1B and Table 1). GUK1 is much faster with GMP (control) than with AT-8001 (respectively 6800 ± 300 nmol/min/nmol protein vs. 34 ± 5 nmol/min/nmol protein). GUK1 exhibits the highest turnover in the activation chain (Table 1), but the 2’-C-methyl, 2’- F modification slows it down ∼200-fold.

GUK1 was crystallized and its structure solved at 1.76 Å resolution in complex with AT-8001 (Fig. 4, Table 2).

**Figure 4.**
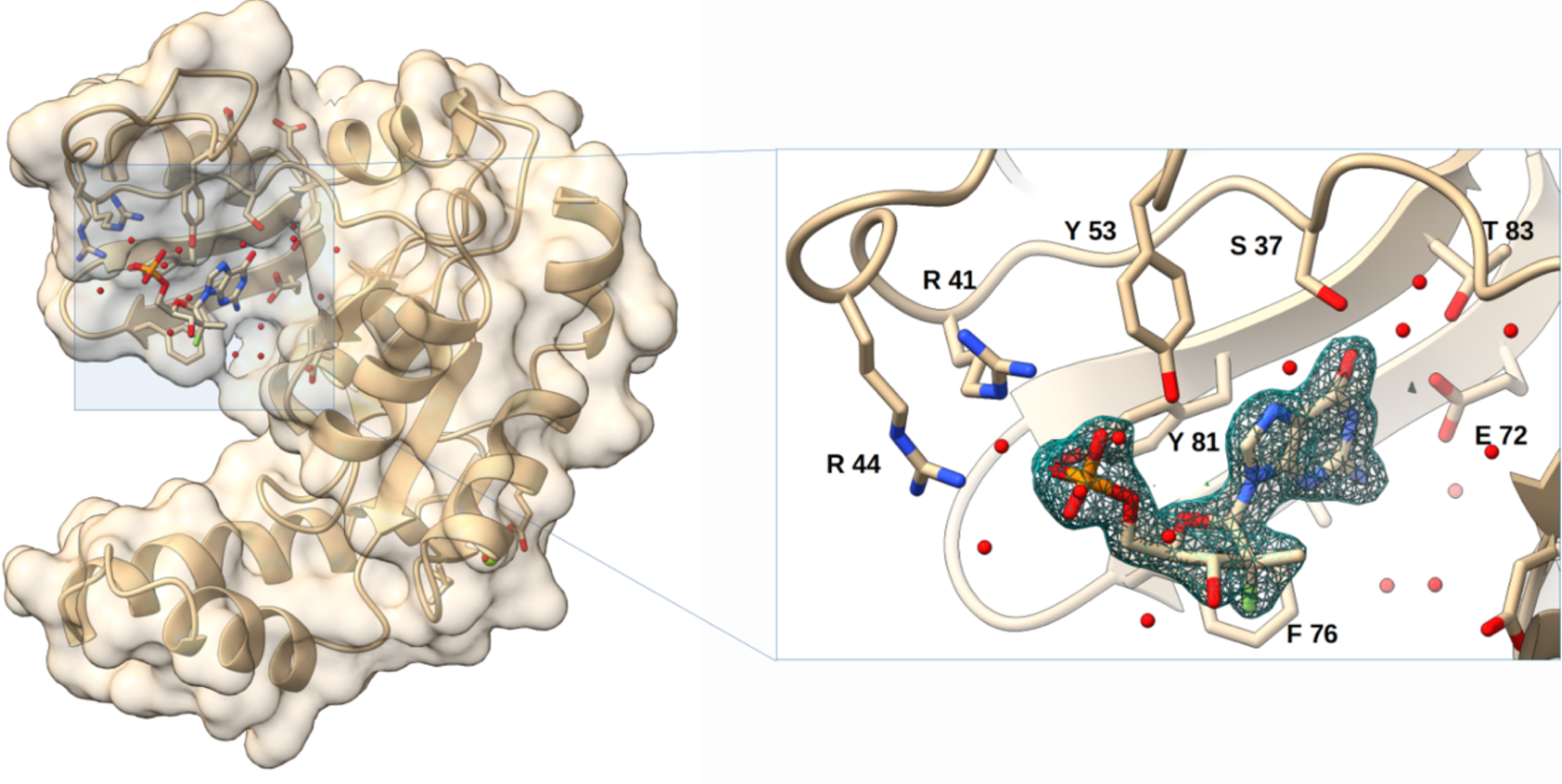
Structure of human GUK1 in complex with AT-8001. Human GUK1 structure is represented in ribbon and transparent surface, while compound is represented in sticks with heteroatom colors. The GUK1 structure corresponds to the open conformation, and molecule is binding to the loading site. On the right side a zoomed part of the molecule is presented in the loading site. Water and side chains of interacting residues with the molecule are shown in sticks. The *F_o_ - F_c_* omit map of the molecule is presented at 1 σ.

**Table 2:** Data-collection and refinement statistics for native human enzymes of the bemnifosbuvir activation pathway complexed with substrate or product compounds.

|  | HINT 1 complexed with<br>AT-8003<br>- 8PWK | ADALP 1 complexed with<br>AT-8001<br>- 8QCH | GUK 1 complexed with<br>AT-8001<br>- 8PTS | NDPK complexed<br>with AT-9010<br>- 8PIE |
| --- | --- | --- | --- | --- |
| Wavelength (Å) | 0.97856 | 0.980 | 0.8856 | 0.978 |
| Resolution range | 39,8 – 2,09<br>(2,13- 2.09) | 105.53 – 2.44<br>(2.48 – 2.44) | 62.85 – 1.76<br>(1.79 – 1.76) | 49.68 – 1.90<br>(1.97 - 1.9) |
| Space group | C2 | P 21212 | C 2 | P21 |
| Unit cell |  |  |  |  |
| a,b,c | 77.96, 46.45, 63.77 | 142.7, 142.9, 156.4 | 132.7, 54.3, 58.9 | 54.2, 120.3, 71.9 |
| $\alpha,\beta,\gamma$ | 90.00, 94.98, 90.00 | 90.0, 90.0, 90.0 | 90.0, 108.8, 90.0 | 90.0, 110.2, 90.0 |
| Total reflections | 89090 | 1216890 | 158084 | 480674 |
| Unique reflections | 12964 | 119069 | 37848 | 67263 |
| Multiplicity | 6.9 (6.4) | 10.2 (10.5) | 4.2 (4.3) | 7.1 (7.2) |
| Completeness (%) | 95.8 (96.8) | 100 (100) | 96 (96) | 98.89 (98.00) |
| Mean I/sigma(I) | 10.6 (3.3) | 7,2 (1,0) | 9.7 (2.2) | 11.03 (1.21) |
| Wilson B-factor | 23,3 | 40,6 | 27.8 | 21.5 |
| R-merge | 0.116 (0.545) | 0.363 (3.973) | 0.055 (0.69) | 0.2401 (0.7668) |
| R-meas | 0.125 (0.592) | 0.383 (4.180) | 0.063 (0.79) | 0.2596 (0.8257) |
| CC1/2 | 0.995 (0.904) | 0.988(0.350) | 0.99 (0.82) | 0.958 (0.745) |
| No. of reflexions<br>used in refinement<br>(free reflections) | 12960<br>(646) | 118763<br>(5715) | 364600<br>(1823) | 67261<br>(6623) |
| R-work | 0,19 | 0.21 | 0.24 | 0,17 |
| R-free | 0.23 | 0.24 | 0.26 | 0,21 |
| Total atoms in<br>structure | 1929 | 23207 | 3232 | 7869 |
| macromolecules | 1756 | 44365 | 3019 | 7287 |
| Ligands | 44 | 320 | 80 | 180 |
| RMS (bonds) | 0.007 | 0.0151 | 0.008 | 0.030 |
| RMS (angles) | 0,095 | 2.253 | 1.86 | 2.28 |
| Ramachandran<br>favoured (%) | 99.55 | 97.27 | 98,4 | 97.95 |
| allowed (%) | 1.06 | 2.0 | 1,6 | 1,4 |
| outliers (%) * | 0.0 | 0.69 | 0.0 | 0.68 |
| Average B-factor | 48.60 | 50.00 | 34.0 | 26.16 |
| Beamline | Soleil Proxima-1 | Soleil Proxima 2 | ESRF ID23-1 | Soleil Proxima-2 |

Statistics for the highest-resolution shell are shown in parentheses.
\*Outliers are in highly flexible regions of the protein.

The overall fold of GUK 1 closely resembles that of the yeast and mouse guanylate kinases^34,35^. Two β-sheets and eight α-helices, the last two separated by a π-helix, define three structurally and functionally distinct sub-domains: the core, the NMP-binding site, and the lid^35^. The core is defined by residues 5–31, 97–123, and 165–194 (helices α1, α4, α7, and α8; strands β1, β7, β8, and β9). The NMP-binding site is a mobile structural element defined by residues 37–89 (starting with a protruding loop and followed by helices α2 and α3; strands β3, β4, β5, and β6). The lid where the phosphate donor (*eg*., ATP) binds, is also mobile and defined by residues 126–156 (helices α5 and α6). These parts are interconnected with four hinges that allow conformational changes from an open to closed state^34^. The structure presented here is an open conformation (Fig. 4), with the AT-8001 substrate binding through extensive contact with both the phosphate moiety and the nucleobase. Thus, on the one hand, Tyr 81, Tyr 53 and Arg 41 share hydrogen bonding with the oxygen of the phosphate, while Arg 44 makes a contact through a water molecule. On the other hand, Glu 72 and Ser 37 engage hydrogen bonds with the base while Tyr 53 contributes to positioning the base through π-stacking. The protein structurally checks the substrate by contacting both the purine base and the presence of a monophosphate, with no significant check on the ribose at this stage.

We modeled AT-8001 into a closed GUK1 conformation based on an energy-minimized model of hGUK1 (Fig. S7). The base is constrained by interactions with Ser 37, Glu 72, and Thr 83. The major difference observed with bound NMP substrates is the ribose pucker conformation. Again, in the structure with bound GMP, the 3’-OH ribose is in *S* pucker position to accommodate the closed conformation of the active site, but not for AT-8001, as described for HINT1 above. To avoid a steric conflict of the 2’-C-methyl, the compound has to slide into the core where the base is π-stacked by Tyr 81 and stabilized by Ser 37 and Glu 72. Both Arg 41 and Arg 44 contact the phosphate; Arg 137 and Arg 148 are expected to catalyze phosphate transfer from ATP bound in the lid, but the repositioning creates distances not compatible for optimal transfer. This positional adjustment likely explains the ∼200-fold discrimination of AT-8001 versus the natural substrate GMP (Table 1).

### Structural and functional basis of NDPK specificity

NDPK is able to transfer a phosphate group from a triphosphate nucleoside donor to a nucleoside diphosphate acceptor^26,36^, to yield the active triphosphate form of bemnifosbuvir AT-9010. The reaction proceeds through a so called ‘ping-pong’ mechanism in which a triphosphate nucleoside transfers covalently its γ-phosphate to the enzyme, and the diphosphate nucleoside acceptor is subsequently bound to the same site to accept the phosphate. NDPK is known to phosphorylate a wide range of nucleotide analogues bearing various structural modifications. The activity of purified NDPK chain b (NDPKb)^27^,is able to phosphorylate GDP (control) into GTP and also AT-8500 into AT-9010, the expected product (Fig. 1B and Table 1). NDPKb shows a high conversion rate and is only 4-fold slower with AT-8500 as substrate than GDP (1300 ± 100 nmol/min/nmol protein vs. 5400 ± 800, respectively).

We crystallized the NDPKb with AT-9010 (Fig. 5).

**Figure 5.**
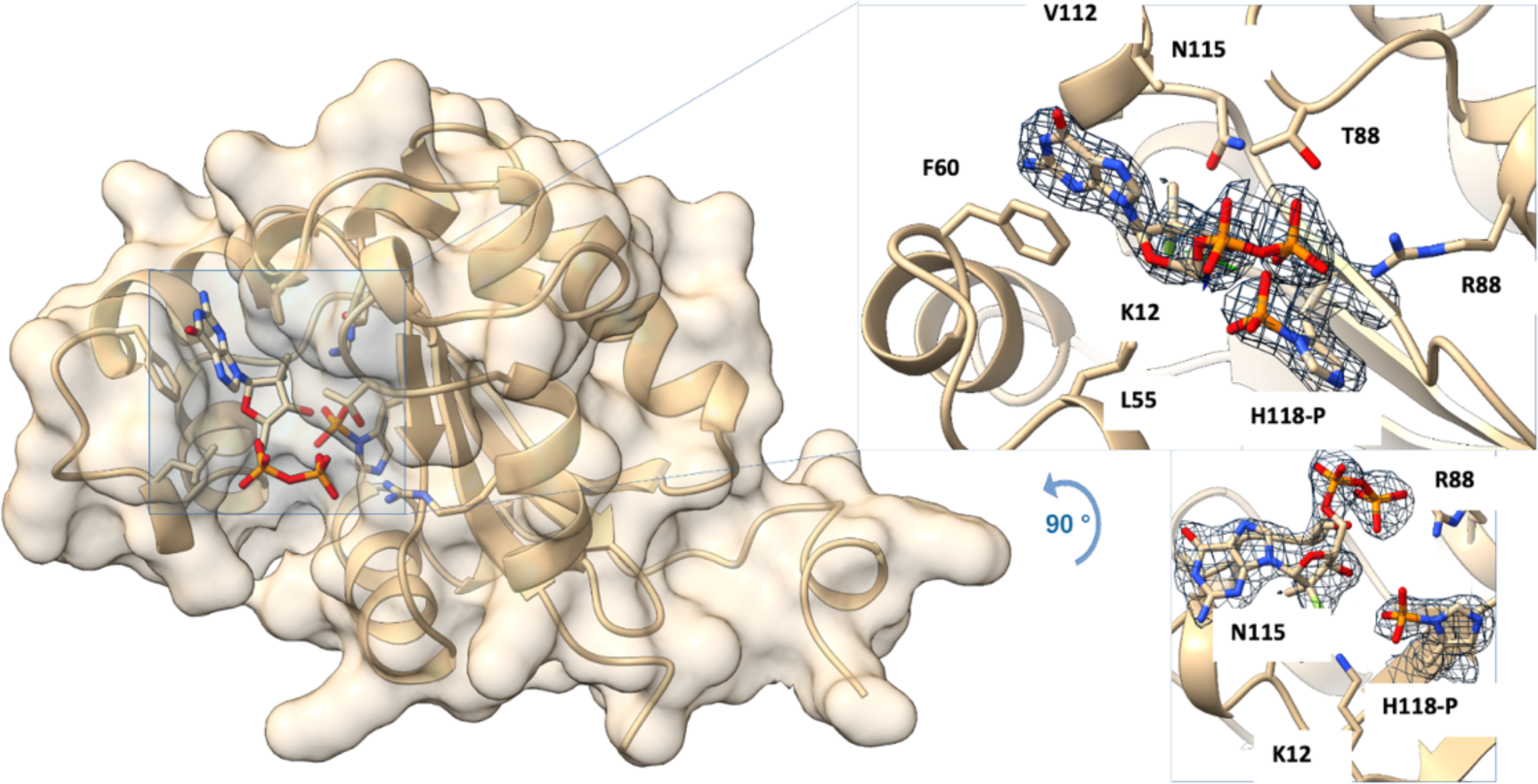
Structure of human NDPKb phosphorylated on H118 and in complex with AT-8500. Human NDPKb structure is represented in ribbon and transparent surface, while compound is represented in sticks with heteroatom colors. On the right side a zoomed part of the molecule is presented in the active site in two orientations differing by 90°. Side chains of interacting residues with the molecule are shown in sticks. The *F_o_ - F_c_* omit map of the molecule as well as the transferred phosphate onto the catalytic histidine is presented at 1 σ.

The crystal structure was solved at 1.9 Å resolution. A hexamer defines the asymmetric unit with six extra densities corresponding to six AT-8500 diphosphate. Interestingly, only the α and β-phosphates of the 5’-triphosphate moiety are visible. A new density is observed near the NE2 of His 118 of each monomer, which correspond to the histidine phosphorylated state His 118- P in Fig. 5 (inset). The AT-9010 γ-phosphate is transferred to the latter due to the protein ability to catalyze the reaction in both directions via a ping-pong mechanism. Structural features of the nucleotide binding site are consistent with the required NDPKb substrate promiscuity. Direct comparison with NDPKb in complex with GDP (PDB: 1NUE) shows a conserved base-stacking between the guanine and Phe 60. The hydrogen bond between Lys 12 and the 3’-OH of the ribose is no longer present because of a displacement of the ribose, probably due to the 2’-C-methyl, 2’-F modification. This shift allows the 3’-OH to make direct contact with the phosphate group of His 118-P (3Å). The β-phosphate is engaged in hydrogen bonds with Arg 88 and is facing the catalytic site of the enzyme.

## Discussion

The journey of a drug begins with cellular uptake up to expression of its antiviral activity into a virus-infected cell. There is a need to better anticipate unexpected complexities. A number of NA prodrugs (*eg*., SOF, TAF, REM) need CatA/CES1^10,11,37,38^-mediated activation. However, CatA is inhibited by the antiviral drug telaprevir (an HCV protease inhibitor)^10,14^, complicating potential combination therapies relying on this esterase. Likewise, the ribavirin antiviral effect results both from cellular inosine monophosphate dehydrogenase and viral growth inhibitions (reviewed in^39^).

Here we cast light on the activation pathway of bemnifosbuvir and its epimer AT-752. Our work quantitates the differential activation of these substrates relative to their natural counterpart GTP. We visualize and measure essential interactions these two compounds and their metabolites engage with specific enzymes involved in their sequential activation.

Turnover measurements (Table 1) show that the rate-limiting steps are generally occurring early in the pathway (CatA/CES1, HINT1, and ADALP1) rather than at the level of the kinases. On these slow, early steps, we note a de-esterification rate ∼14-fold higher for AT-511 than AT-281 into the common intermediate AT-551 by CatA. Comparatively, CES1 reacts ∼300- fold slower than CatA with AT-511, but shows an opposite stereo-preference with AT-281 being preferred ∼8-fold over AT-511. These results are aligned with those obtained with PSI-7851, a racemic mixture of PSI-7976 and PSI-7977 (SOF) which shows both the same CatA and stereoisomer preference^14^.

Murakami *et al.*^14^ have shown that HINT1 is the rate-limiting step along the SOF activation pathway. Its metabolic intermediate PSI-352707 (Fig. S1) binds poorly to HINT1, precluding saturating substrate conditions in infected cells and determination of kcat and Km in enzyme assays. In the case of AT-551, HINT1 is also the rate-limiting step (Table 1). Our HINT1:AT-8003 complex structure provides a structural basis for the kinetic bottleneck along the SOF activation line^14^. We propose that the higher HINT1 conversion activity of AT-551 as compared to the corresponding intermediate metabolite of SOF, PSI-352707, which differ only by their nucleobase, results from the higher stacking power of purines versus uracil onto Ile 44. GS-6620 (Fig. S1) is a C-nucleoside combining a purine-like base with a 2’-hydroxy, 2’-C-methyl and a 1’-CN group. Murakami *et al.* have shown that GS-6620 has a comparable HINT1 turnover to that of AT-551^37^ suggesting that GS-6620 likely adopts the same binding mode as AT-551 rather than fitting into the pocket able to tightly accommodate natural purines. The HINT1 limiting step, however, does not seem to impact accumulation of the SOF 5’- triphosphate form (nor AT-9010) in cells. The compound pools are then readily available for selective use by the NS5B viral RdRp in HCV-infected liver cells.

These results indicate that cell types and their relative expression of these ‘early step’ enzymes CatA/CES1 and HINT1 are expected to play a major role in drug activation: *i*) the esterase-mediated deprotection step may well govern the overall rate of conversion of the drug into its active metabolite; *ii*) the relative abundance of CatA- and CES1-like enzymes should influence these early steps; and *iii)* further optimization in antiviral activity might be possible through optimization of the prodrug part in the 5’ portion of the NA rather than focusing on optimizing the kinase steps. The choice of targeted infected cells may ultimately point to the most appropriate epimer and prodrug. In the case of ADALP1, unlike HINT1, the substrate and products are superimposable in the crystal structure, indicating that this complex should be reliable and useful for drug design.

Ideally, the NA activation line must be proven functional in cells, organs, or animal models supporting pre-clinical tests before commitment to full clinical trials in humans. AlphaFold2^40^- generated structural models of homologous animal enzymes did not point to obvious polymorphisms potentially deleterious for activation (not shown). The very high degree of structural conservation along species suggests a vital role in mammals for these activation enzymes. As such, monitoring differential expression of HINT1 should remain a sufficient and essential asset in predicting P-N hydrolysis in a given tissue or cell. The situation is somewhat similar for ADALP1, which shows significant polymorphisms across organisms, although mapping them invariably points to external, solvent exposed loops, with no direct impact on the NA binding site.

In the case of exposition of cells to bemnifosbuvir, its corresponding 5’-triphosphate AT-9010 has been demonstrated in cellular concentrations up to ∼700 µM in human primary cells and cell types incubated *in vitro* with AT-511, including bronchial and nasal epithelial cells^18^, hepatocytes and Huh7 cells^24^, and peripheral blood mononuclear cells (data not shown). Thus, none of the enzymes studied here limit the formation of the active compound AT-9010 in these cells. Bemnifosbuvir, though, shows reduced antiviral activity in some cell lines, *eg.*, in VeroE6, HeLa, or MRC-5 cells^18^. We note that *cynomolgus* monkey hepatocytes have been reported to activate bemnifosbuvir to levels 50-fold lower than Huh7 cells^18^. The superimposition of human and *cynomolgus* HINT1 and ADALP1 structures does not point to any polymorphism that could substantiate differences in substrate binding or catalysis (not shown).

Bemnifosbuvir shares two activation enzymes (HINT1 and NDPKb) with SOF and RDV. Hence, the HINT1 structure should represent the most relevant enzyme to optimize the prodrug part of such antivirals. Two other enzymes ADALP1 and GUK1 are specific for diamino purine- and guanine-containing NAs, respectively. With the abacavir success in mind^22^, the ADALP1 structure presented here should guide further exploration of chemical diversity at the *N^6^*-amino purine group.

A wealth of structural data already exists regarding nucleotide analogues in complex with RNA virus replicases, *eg*., HCV^15^; picornavirus^41^; Reovirus^42^; SARS-CoV-2^13,28,43^; for a repository of stacked RdRp structures, see^44^. Much like along the NA activation line, analysis of selectivity of viral polymerase in ternary complexes also identifies proximal amino acid side chains potentially able to accommodate chemical decorations without impacting the NA-5’- triphophate incorporation efficiency.

Finally, a NA drug potential is also conversely determined by its innocuity towards host enzymes that could misuse their 5’-triphosphate form and alter cell metabolism or host nucleic acid. Human mitochondrial RNA polymerase (POLRMT) inhibition accounts, in part, for cellular toxicity of NAs^45^. Comparative examination of ternary complexes of viral RdRps and POLRMT^46^ with their RNA and NA-TP substrates may further document the strategy to rationally integrate NA drug design from chemistry to selective inhibition of viral growth^45^.

Our work casts light on a whole set of previously ill-defined reactions involved in drug activation, and contributes to the integration of nucleotide analogue drug-design from bioavailability to precise and selective mechanism of action as direct-acting antivirals.

## Materials and Methods

### Materials

Recombinant human cathepsin A (CatA), cathepsin L (CatL), carboxylesterase 1 (CES1) and *trans*-epoxy-succinyl-_L_-leucylamino(4-guanidino)butane (E-64) were purchased from R&D Systems. The respective genes coding for the human recombinant enzymes Histidine triad nucleotide binding protein 1 (HINT1), Adenosine deaminase-like protein 1 (ADALP1), Guanylate kinase 1 (GUK1) and Nucleoside diphosphate kinase (NDPKB) were coded into a pNC-ET28 expression vector with an N-terminal His-tag and TEV cleavage site, purchased from Twist Bioscience. Control compound AMP-NH_2_ was from Biosynth, *N^6^*-Me-AMP was purchased from BioLog Life Science Institute and AMP, IMP, GMP, GDP and GTP were from Sigma Aldrich. Other compounds with a name starting with “AT” were synthesized by TopPharm, Inc., USA.

### Protein expression

The DNA sequence encoding for Adenosine deaminase-like protein 1 (ADALP1), Nucleoside diphosphate kinase B (NDPKB), Guanylate kinase 1 (GUK1) and Histidine triad nucleotide binding protein 1 (HINT1) were synthesized and cloned into a pET-28a (+) vector for HINT1 and GUK1, into a pNC-ET28 vector for NDPKB and into a pET-28 for ADALP1 (Twist Biosciences). The final construct encoded for HINT1, ADALP1, GUK1 and NDPKB with a N-terminal hexahistidine tag (6xHis) and a TEV protease recognition site.

All proteins were expressed in *E. coli* NEB C2566 cells (New England Biolabs). The cells were grown at 37°C in TB medium for HINT1, GUK1 and NDPKB, LB Borth for ADALP1 containing 50µg/mL Kanamycin until the absorbance at 600nm reached 0.6 – 0.8. At this stage induction was started with 0.5mM IPTG and the cells were grown overnight at 16°C. Then the cells were harvested and the pellets were suspended in lysis buffer (50mM Tris pH = 8.0, 300mM NaCl, 10mM imidazole, 1mM PMSF, 0.25mg/mL lysozyme and 10µg/mL DNase) for HINT1, GUK1 and NDPKB, before storing them at −80°C. For ADALP1 the pellet was freeze-dried and stored at -80°C.

### HINT1, GUK1 and NDPKb purification

The lysate was disrupted and cleared by centrifugation at 12,000g for 30min at 10°C. Then the supernatant was loaded onto a Ni-NTA-beads (ThermoFisher) batch and washed with 50 mM Tris pH = 8.0, 300 mM NaCl, 20 mM imidazole buffer. The recombinant protein was then eluted with 50 mM Tris pH = 8.0, 300 mM NaCl, 250 mM imidazole buffer. 10% glycerol was added for HINT1. TEV protease was used to remove the N-terminus His-tag during an overnight dialysis in a buffer (50 mM Tris pH = 8.0, 150 mM NaCl, 1 mM DTT) at 4°C for HINT1, at 37°C for GUK1and at room temperature for NDPKB.

A second purification step on Ni-NTA beads was performed to remove any uncleaved protein before to achieve a SEC purification onto a Superdex 75 16/600 GE column (GE Healthcare) in a final buffer containing 10 mM Tris pH = 8.0, 50 mM NaCl for HINT1, 20 mM Tris pH = 8.0, 100 mM NaCl, 1 mM DTT for GUK1 and 50 mM Tris pH = 8.0, 150 mM NaCl, 1 mM DTT for NDPKb. The fractions were analysed on SDS-PAGE and the ones containing the pure target protein were pooled. The purified protein was then concentrated until 10 mg/mL, aliquoted and stored at -80°C. Same protocol was used to express and purify uncleaved NDPKb for activity assays.

### ADALP1 purification

Bacterial pellets were lysed in half buffer (50 mM HEPES pH = 7.5, 300 mM NaCl, 10% glycerol, 0.5 mM TCEP) and half Master Mix Bug Buster (Merck) supplemented with a tablet of EDTA-free antiprotease cocktail (Roche) per 50 ml of lysate before sonication.

The lysate was cleared by centrifugation at 12,000g for 30min at 10°C and the supernatant was applied onto a Talon Superflow beads batch (Merck). The immobilized proteins were washed with 50 mM HEPES pH 7.5, 300 mM NaCl, 12.5 mM Imidazole, 10% glycerol, 0.5 mM TCEP, then washed with 50 mM HEPES pH 7.5, 1.5M NaCl, 10% glycerol, 0.5 mM TCEP and eluted in buffer (50 mM HEPES pH 7.5, 300 mM NaCl, 150 mM Imidazole, 10% glycerol 0.5 mM TCEP). TEV protease was used to remove the N-terminus His-tag during an overnight dialysis in a buffer 50 mM HEPES pH = 7.5, 300mM NaCl, 10% glycerol, 1mM DTT. The untagged ADALP1 was further purified by a Talon Superflow beads batch (Merck).

Finally, ADALP1 was purified by size exclusion chromatography using a S75 16/60 GE Column (GE Healthcare) equilibrated in buffer containing 10 mM Tris pH = 8.0, 40 mM NaCl, 1 mM TCEP. The single ADALP1 peak corresponded to the monomeric form. It was concentrated to 10 mg/mL, flash-frozen in liquid nitrogen, and stored at −80°C.

### CatA enzyme assay

Human recombinant CatA was first activated by following the manufacturer’s instructions (R&D Systems), incubating at 37°C 10 µg/mL CatA with 1 µg/mL CatL in an activation buffer (25 mM MES pH = 6.0, 5 mM DTT) during 30 min. 10 µM E-64 (CatL inhibitor) was then added before aliquoting and storage of activated CatA at -80°C. AT-511 and AT-281 hydrolysis by activated CatA was measured by incubating 100 µM compound in a reaction buffer containing 25 mM MES pH 6.5, 100 mM NaCl, 1 mM DTT, 0.1% NP-40 and 20 nM enzyme at 37°C for 45 min. The reaction was started by adding the enzyme. At various time points, 10 µL aliquots were collected from the reaction mixtures mixed with EDTA 10 mM (final concentration) and stopped by heating the samples at 95°C for 5 min. The sample were filtered on centrifugal filters Microcon® - 10 (Sigma Aldrich). The filtrates were mixed with TEAB 1M (1:1) and injected onto a C18 reverse phase column (2.5 µm, 4.6 by 100 mm, Xbridge C18 BEH, Waters) equipped with a guard column, equilibrated with 98:2 TEAB 50 mM pH = 7 (buffer A):MeCN. The substrates and reaction products were eluted using a non-linear gradient of acetonitrile in buffer A, detailed in supplementary data.

### CES1 enzyme assay

AT-511 and AT-281 hydrolysis by human recombinant CES1 (100 nM) was assayed by incubating at 37°C the enzyme with 100 µM compound in 50 mM Tris buffer pH = 7.5, 0.1% NP-40 and 1 mM DTT for 2h. The reaction was started by adding the substrate. At various time points, 20 µL aliquots were collected from the reaction mixtures mixed with EDTA 10 mM (final concentration) and stopped by heating the samples at 95°C for 5 min. The sample were filtered on centrifugal filters Microcon® - 10 (Sigma – Aldrich). The filtrates were mixed with TEAB 1M (1:1) and injected onto a C18 reverse phase column (2.5 µm, 4.6 by 100 mm, Xbridge C18 BEH, Waters) equipped with a guard column, equilibrated with TEAB 50 mM pH = 7 (buffer A). The substrates and reaction products were eluted using a non-linear gradient of acetonitrile in buffer A, detailed in supplementary data.

### HINT1 enzyme assay

The hydrolytic reactions with HINT1 (100 nM) were performed on AMP-NH_2_ and AT-551 (200 µM) in 20 mM HEPES buffer pH 7.2, 20 mM KCl, 1 mM MgCl_2_ and 1 mM DTT, at 37°C for 2h. The reaction was started by adding the enzyme. At various time points, 10 µL aliquots were collected from the reaction mixtures mixed with EDTA 10 mM (final concentration) and stopped by heating the samples at 95°C for 5 min. The sample are filtered on AcroPrep Advance 96-Well Filter Plates with 3K Omega membrane (Pall). The filtrates were mixed with TEAB 1M and injected onto a C18 reverse phase column (3 µm, 3 by 150 mm, Acclaim Polar Advantage II, ThermoFischer) equipped with a guard column, equilibrated with TEAB 50 mM pH = 7 (buffer A). The substrates and reaction products were eluted using a non-linear gradient of acetonitrile in buffer A, detailed in supplementary data.

### ADALP1 enzyme assay

ADALP1 (100 nM) activity was measured on several substrates (*N^6^*-Me-AMP, AT-8003, AT-8002 AT-8004, AT-551, AT-259, AT-229) at 200 µM concentration in a reaction buffer containing BTP pH = 6.8, 100 mM NaCl and 1 mM DTT. Reactions were incubated at 37°C for 2h. 1 µM ADALP1 was also tested with AT-8002, AT-551, AT-229 and AT-259. The reaction was started by adding the substrate. At various time points, 10 µL aliquots were collected from the reaction mixtures diluted with H_2_O and stopped by heating the samples at 95°C for 5 min. The sample were filtered on AcroPrep Advance 96-Well Filter Plates with 3K Omega membrane (Pall). The filtrates were mixed with TEAB 1M and injected onto a C18 reverse phase column (2.5 µm, 4.6 by 100 mm, Xbridge C18 BEH Premier, Waters) equipped with a guard column, equilibrated with TEAB 50 mM pH = 7 (buffer A). The substrates and reaction products were eluted using several non-linear gradients of acetonitrile in buffer A, detailed in supplementary data.

### GUK1 enzyme assay

GUK1 (40 nM) was assayed on GMP and AT-8001 (200 µM) in a reaction buffer containing 50 mM Tris buffer pH = 8.0, 50 mM KCl, 5 mM MgCl_2_, 1 mM ATP and 1 mM DTT. Reactions were incubated at 37°C for 1h. The reaction was started by adding the enzyme. At various time points, 10 µL aliquots were collected from the reaction mixtures, mixed with EDTA 10 mM (final concentration) and stopped by heating the samples at 95°C for 5 min. The sample were filtered on AcroPrep Advance 96-Well Filter Plates with 3K Omega membrane (Pall). The filtrates were mixed with TEAB 1M and injected onto a C18 reverse phase column (3 µm, 3 by 150 mm, Acclaim Polar Advantage II, ThermoFischer) equipped with a guard column, equilibrated with TEAB 50 mM pH = 7 (buffer A). The substrates and reaction products were eluted using a non-linear gradient of acetonitrile in buffer A, detailed in supplementary data.

### NDPKB enzyme assay

NDPKB (20 nM) was assayed on GDP and AT-8500 (200 µM) in a reaction buffer containing 50 mM Tris buffer pH = 8.0, 50 mM KCl, 5 mM MgCl_2_, 1 mM ATP and 1 mM DTT. Reactions were incubated at 37°C for 1h. The reaction was started by adding the enzyme. At various time points, 10 µL aliquots were collected from the reaction mixtures mixed with EDTA 10 mM and stopped by heating the samples at 95°C for 5 min. The sample were filtered on AcroPrep Advance 96-Well Filter Plates with 3K Omega membrane (Pall). The filtrates were mixed with TEAB 1M and injected onto a C18 reverse phase column (3 µm, 3 by 150 mm, Acclaim Polar Advantage II, ThermoFischer) equipped with a guard column, equilibrated with TEAB 50 mM pH = 7 (buffer A). The substrates and reaction products were eluted using a non-linear gradient of acetonitrile in buffer A, detailed in supplementary data.

### Crystallization of HINT1

Crystallization conditions were adapted from^21^. Briefly, co-crystals with AT-8003 were grown at 293.15 K, using a 1:1 ratio of HINT1 at 10 mg/mL with AT-8003 (final concentration 25 mM) to precipitant solution 0.1M MES pH 6.1-6.5, 27%-30% PEG 8000. Crystals grew in a few days and were cryo-protected with reservoir solution supplemented with 20% PEG 400, and flash-frozen in liquid nitrogen at 100 K.

### Crystallization of ADALP1

Crystallization assay were set up using the sitting drop vapor diffusion method, using a 2:1 ratio of protein solution to the reservoir solution at 293.15K. The co-crystals with AT-8001 were obtained by mixing ADALP1 at 17 mg/mL with AT-8003 (final concentration 10 mM) and 1:10000 w/w dilution a-Chymotrypsin prior to crystallization and were grown from 1.1-2.1 M ammonium sulfate, 0.1 M sodium cacodylate/HCl pH 5.3-6.3, 0.2 M sodium chloride for 2 months. Crystals were cryo-protected with reservoir solution supplemented with 20% glycerol, and flash-frozen in liquid nitrogen at 100 K.

### Crystallization of GUK1

Large co-crystals of GUK1 (17 mg/mL) with AT-8001 (2 mM) supplemented with 5 mM MgCl2 were grown using the sitting drop method with a 3:1 ratio of protein solution to precipitant solution consisting of 66 mM Sodium Cacodylate pH 6.5-7.5, 12.2-22.2% PEG 3350, 8.3% PEG 4000, 33 mM MES pH6.5 and 66 mM magnesium chloride over a period of 10 days at 293.15K. Crystals were cryo-protected with reservoir solution supplemented with 20% glycerol, and flash-frozen in liquid nitrogen at 100 K.

### Crystallization of NDPKb

Crystallization conditions were adapted from^47^ using the sitting-drop vapor diffusion method using crystallization buffer. All crystals were grown at 277.15K, using a 1:1 ratio of protein (5 mg/mL) mixed with AT-9010 (final concentration 1 mM) to precipitant solution (12% PEG 4000, 50 mM Tris pH = 8.4, 16% glycerol, 1 mM DTT). Crystals grew in 3 days and were flash-frozen directly in liquid nitrogen at 100 K.

### Structure Determination of the Human HINT 1: AT-8003 complex

The dataset of HINT1 in complex with AT-8003 was collected on the Proxima-1 beamline at the Synchrotron SOLEIL. Dataset was processed using AUTOPROC^48^. The phase was obtained using Molecular Replacement using MOLREP^49^ with the PDB entry 6N3V as a model. The ligand AT-8003 and geometry description files were generated using the GRADE2 server^50^. Structure handling and refinement were done using COOT^51^ and BUSTER^52^.

### Structure Determination of the Human ADALP 1 : AT-8001 complex

The dataset of ADALP 1 in complex with AT-8001 was collected on the Proxima-2 beamline at the Synchrotron SOLEIL. Dataset was processed using AUTOPROC^48^ and processed data were sent to the CCP4 cloud suite^53^. The pipeline is fully automatic. The phase was obtained using Molecular Replacement – MORDA^53^ using PDB entry 6IV5 as a model. The ligand AT-8001 and geometry description files were generated using the GRADE2 server^50^. Minor correction on the structure and further refinement were done using COOT^51^ and REFMAC5^54^, respectively.

### Structure Determination of the Human GUK 1 : AT-8001 complex

The dataset of GUK1 in complex with AT-8001 was collected on the ESRF ID23-1 beamline. Dataset was processed using AUTOPROC^48^. The phase was obtained using Molecular Replacement – MORDA^53^ using PDB entry 4F4J as a search model and optimized with ARP/WARP^55^. Structure refinement was done using COOT^51^ and BUSTER^52^, respectively.

### Structure Determination of the Human NDPK : AT-8500 complex

The dataset of hNDPKb in complex with AT-8500 was collected on the Proxima-2 beamline at the Synchrotron SOLEIL. Dataset was handled using the CCP4 suite^56^. Images were processed by XDS^57^ and AIMLESS^58^. The phase was obtained using Molecular Replacement – PHASER^59^ with the PDB entry 1NUE as a model. The ligand AT-8500 corresponding to the diphosphate form of AT-9010 was generated using the AceDRG program^60^. Structure handling and refinement were done using COOT^51^ and REFMAC5^54^software, respectively.

All electron-density maps were inspected using COOT^51^. Extra density accounting for ions, and/or compounds were observed for all complexed structures. The structures were evaluated using MOLPROBITY^61^ and PROCHECK^62^. Structural analysis and high resolution figures were done with UCSF ChimeraX^63^. Facilities, statistics of data collections, refinements and PDB deposition code are given in Table 1.

### Modeling of GUK 1 closed conformation

Sequence of human GUK 1 was submitted to an homology modeling process using modeler V10 and the structure of mouse GUK complexed with GMP and ADP (PDB 1LVG) as template. The two sequence have 88 % identity, which makes us confident that the resulting closed conformation model is reasonable.

### Docking of GMP and AT-8001 in hGUK 1 closed conformation

The 3D model of hGUK 1 was energy minimized by the steepest gradient method of energy minimization followed by conjugate gradient minimization, using the MMTK63 and AMBER64 packages. Mol2 and PDB files format of the ligands and receptor were converted to PDBQT format using chimera prior to docking. All the water and solvent atoms of the protein were removed and the polar hydrogen and polar charge were added onto the ions and ligand prior to docking. The protein was kept rigid while the ligand was allowed to rotate and explore more flexible binding pockets. Docking of the respective ligands into the cavity were performed iteratively using AUTODOCK VINA65. The best poses from the first round of docking were used as seed for the second round. The resulting first round of docking were carefully analyzed to retained the best poses. The grid box size dimensions were first 40X40X40, to verify that our ligands will preferentially bind in the catalytic site. The grid box size was further optimized to 23.2X25.6X21.2 thus covering the binding pockets, the default scoring function was used for docking. As control for the procedure GMP was docked following the same protocol and the final pose is virtually identical to the one measure in the experimental structure (PDB 1LVG). Binding modes of the docked complexes were obtained and sorted based on their binding energy, ions and amino acid residues present at a distance less of 3 Å were considered as the binding partners of the ligands. Binding modes were compared to theses of the native structure. The interaction figures representing the docked complexes have been generated using ChimeraX^63^.

## Acknowledgments

We thank Veronique Fattorini, Camille Falcou, and Pierre Gauffre for synthetic gene design and help in protein purification. We thank Nancy Agrawal, Alex Vo, and Qi Huang for critical reading of the manuscript, together with Ashleigh Shannon and numerous colleagues for helpful suggestions along the project.

The authors acknowledge SOLEIL and ESRF for provision of synchrotron radiation facilities and we would like to thank beamline teams of Proxima-1, Proxima-2 and ID23-eh1 for their assistance during data collection. This work was supported by the ‘Agence Nationale pour la Recherche’ through the French Infrastructure for Integrated Structural Biology (FRISBI) [ANR-10-INSB-05-01].

## Author contributions

Conceptualization: KA, FF, BC

Methodology: KA, FF, BC

Experiments: AC, CZ, MF, AM, FF

Analysis: AC, MF, AM, SG, JPS, KA, FF, BC

Funding acquisition: BC Writing: FF, BC

## Data and materials availability

The coordinates and structure factors for HINT1, ADALP1, GUK1, and NDPKb structures have been deposited in the PDB with accession codes 8PWK, 8QCH, 8PTS, and 8PIE, respectively.

