## Supplementary material for "The activation chain of the broad-spectrum antiviral bemnifosbuvir at atomic resolution"

### Supplementary Figures

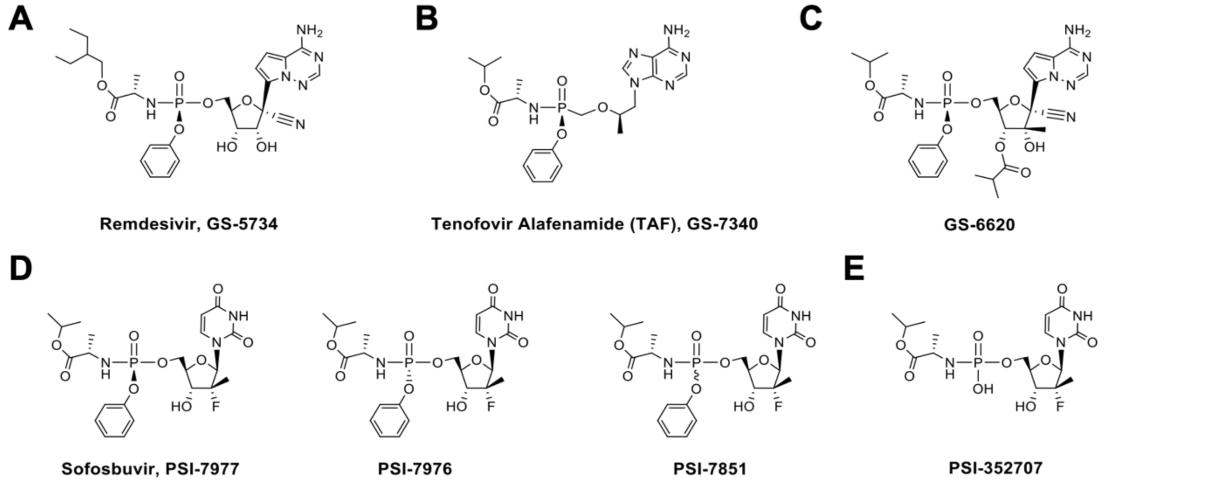


**Fig. S1. Structure of NAs other than-AT-compounds.** (A) Remdesivir or GS-5734, aryloxy phophoramidate prodrug of an adenosine analogue, with 1′ -C-nucleoside bond and 1’-cyano substitution. (B) Tenofovir Alafenamide (TAF) or GS-7340, prodrug of tenofovir, acyclic phosphonate analog of adenosine monophosphate. (C) GS-6620, aryloxy phosphoramidate prodrug of an adenosine analogue, with 1’-C-nucleoside bond and 1’-cyano-2’-C-methyl substitutions. (D) Sofosbuvir or PSI-7977 ($S_P$ diastereoisomer) and related compounds PSI-7976 ($R_P$ diastereoisomer) and PSI-7851 (mixture of both diastereoisomers), aryloxy phosphoramidate prodrug of uridine analogue with a 2′-fluoro-2′ -C-methyl modified ribose. (E) PSI-352707, phosphoramidate metabolite of sofosbuvir.

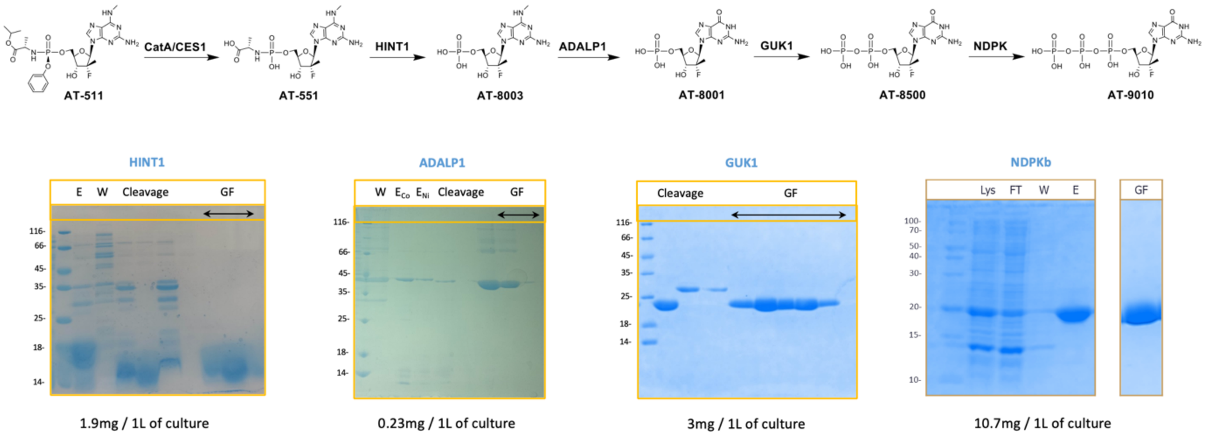


**Fig. S2. Gels SDS-PAGE of purified enzymes HINT1, ADALP1, GUK1 and NDPKb**. Key : Lysate (Lys), Flow-through (FT), Wash (W), Eluate (E), Tag cleavage eluate (Cleavage), Gel filtration (GF).

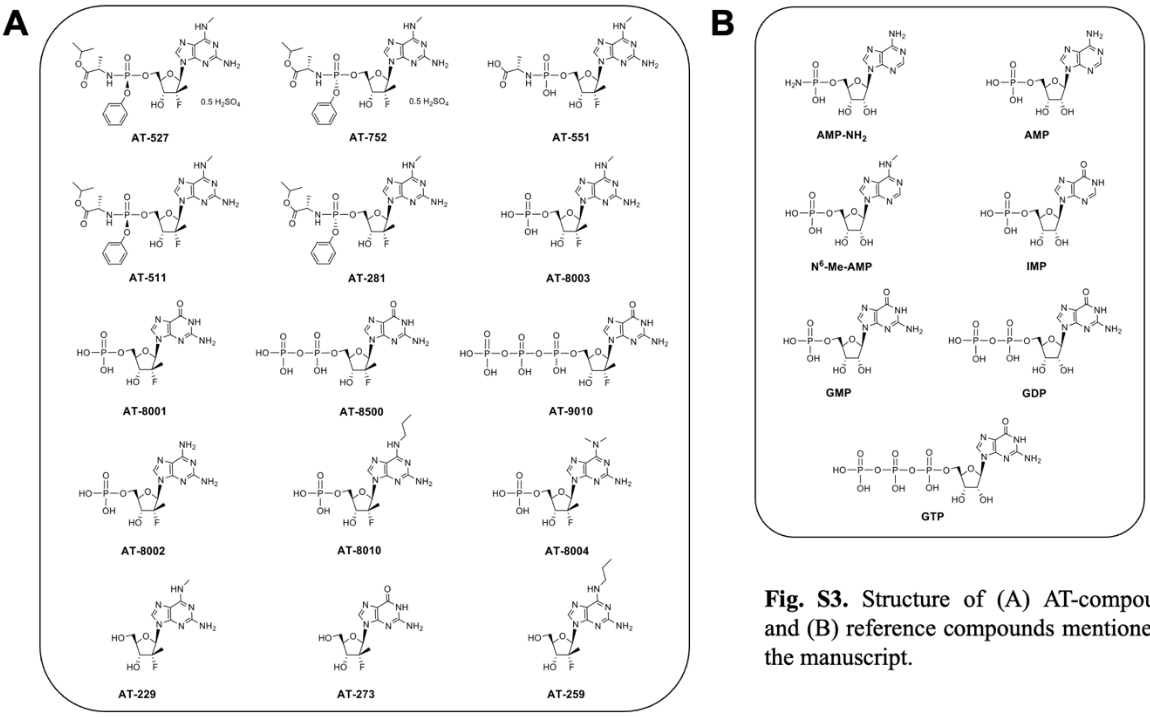


**Fig. S3.** Structure of (A) AT-compounds and (B) reference compounds mentioned in the manuscript.

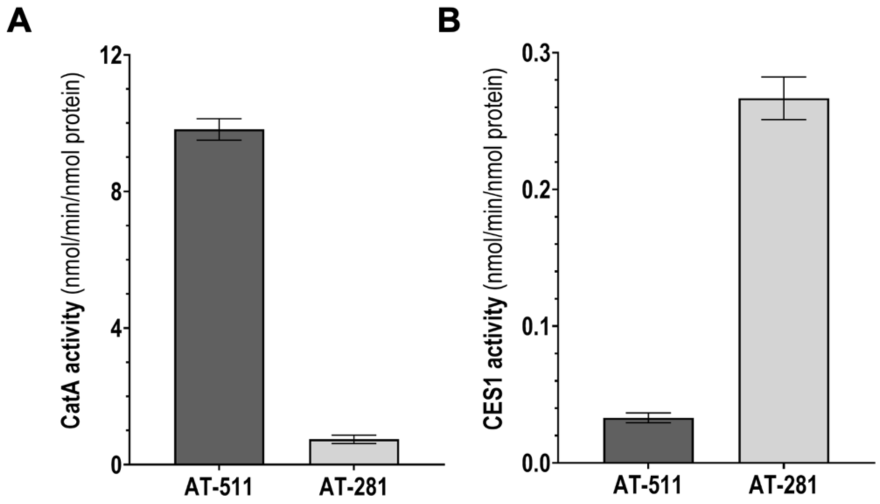


**Fig. S4 : CatA/CES1 stereoselectivity.** Activity of CatA or CES1 with either AT-511 ($S_P$ isomer) or AT-281 ($R_P$ isomer) as substrates. 20 nM CatA was incubated 45 min at 37°C with 100 µM substrate. 100 nM CES1 was incubated 2h at 37° with 100 µM substrate. Bars show mean values (± SD) of three independent experiments.

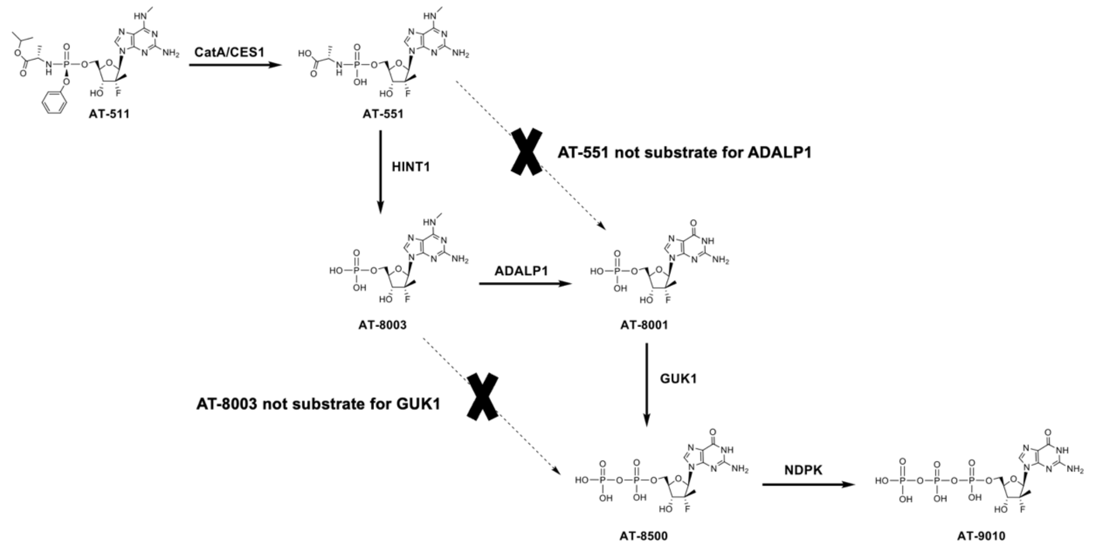


**Fig. S5. Specificity and order of the reactions**. Activation pathway of bemnifosbuvir follows this specific order of reaction. As shown in Table 1, activity assay of ADALP1 with AT-551 as substrate and GUK1 with AT-8003 as substrate did not show any conversion even with 10-fold more enzyme than our standard protocol.

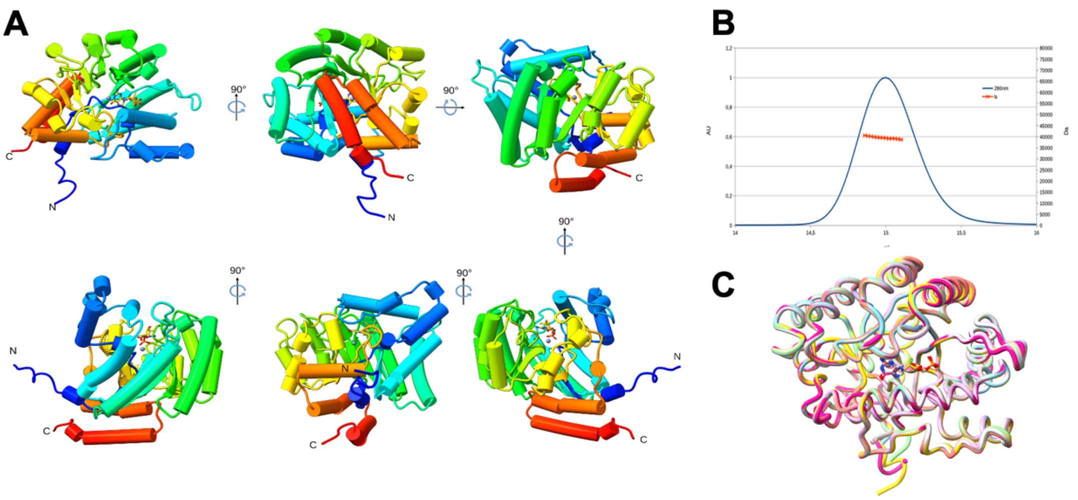


**Fig. S6. Human ADALP1 structure analysis. A)** The protein structure is represented in cylinders and stubs and colors in rainbow color code. Different orientations are presented in respect to the previous one. **B)** Multiple Angle Light Scattering result graph, presenting the elution curve followed at 280 nm (blue line) and the corresponding scattering graph (orange line), experiment show the homogenity of the sample. **C)** The asymmetric unit contained 8 molecules that all contained the compound, presented the superimposition of the 8 chains present showing that they are virtually identical.

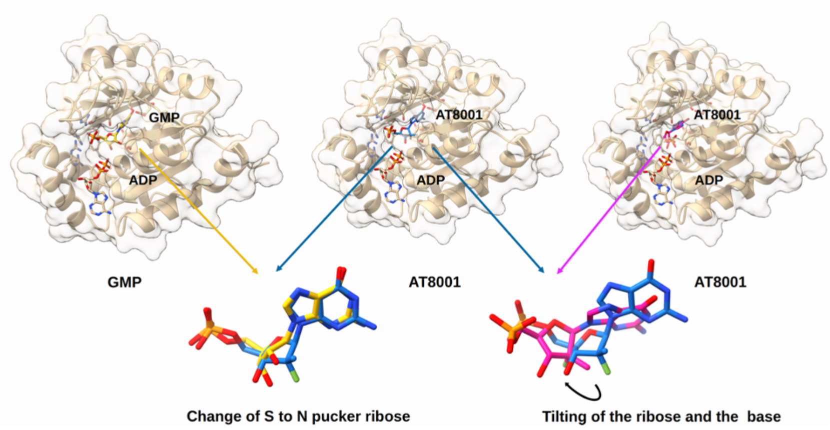


**Fig. S7. A model of human GUK1 in its closed conformation.** The human GUK1 model is represented in ribbon and transparent surface, while the compound is represented in sticks with heteroatom colors and ADP (brown) at the active site. In the left panel is presented GUK1 in complex with GMP (yellow) ; in the central and right panel are presented respectively GUK1 in complex with AT-8001 in its theoretical position (blue) *i.e* superposed with GMP and docked (pink) in the cavity. Below is presented a comparison of the superimposition of the GMP with the theoretical position of AT-8001, highlighting a different ribose pucker conformation ; and the superimposition of the theoretical position of AT-8001 with the docked AT-8001, highlighting a tilting of the ribose and the base to reduce steric hindrance between the 2' methyl with the main chain.
